# Endothelial Baf60c Alleviates Hyperoxia-Induced BPD-Associated Pulmonary Hypertension Through Smarcc2 and PI3K-Akt-mTOR Signaling

**DOI:** 10.64898/2026.06.26.734924

**Authors:** Qiqi Li, Qianyun Cao, Lu Zu, Qiaozhi Wu, Kewei Chen, Chengcheng Hang, Lizhong Du

**Affiliations:** Department of Neonatology, Children’s Hospital, Zhejiang University School of Medicine, National Clinical Research Center for Children and Adolescents’ Health and Diseases, Hangzhou, Zhejiang, China

**Keywords:** bronchopulmonary dysplasia, pulmonary hypertension, pulmonary microvascular endothelial cells, Baf60c, Smarcc2, PI3K-Akt-mTOR, hyperoxia, chromatin remodeling

## Abstract

**BACKGROUND:** Bronchopulmonary dysplasia–associated pulmonary hypertension (BPD-PH) complicates prematurity and carries substantial morbidity in extremely preterm infants. Pulmonary microvascular endothelial cell (PMVEC) dysfunction promotes capillary rarefaction and vascular remodeling, but epigenetic mechanisms after neonatal hyperoxia are poorly defined. Baf60c (SMARCD3), a SWI/SNF subunit supporting vascular homeostasis, and Smarcc2 (BAF170), a PBAF scaffold subunit linked to proliferative signaling, have not been studied together in BPD-PH.

**METHODS:** Neonatal C57BL/6 mice were exposed to 85% oxygen for 14 days. Right ventricular systolic pressure (RVSP), right ventricular hypertrophy, lung weight index, and pulmonary histopathology were assessed; PMVEC proliferation, migration, and invasion were measured. Transcriptome sequencing with GO/KEGG analyses, siRNA knockdown, LY294002 inhibition, coimmunoprecipitation, and Western blotting mapped the Baf60c–Smarcc2–PI3K-Akt-mTOR axis. A Tie1-driven, lung-tropic adeno-associated virus delivered by superficial facial vein injection at postnatal day 1 enabled PMVEC-specific Baf60c overexpression.

**RESULTS:** Hyperoxia increased RVSP, right ventricular hypertrophy, and lung weight index, impaired alveolarization, reduced capillary density, and promoted arteriolar remodeling. PMVEC function was impaired, with PI3K-Akt pathway enrichment and suppressed signaling. Hyperoxia decreased Baf60c and increased Smarcc2. Baf60c knockdown upregulated Smarcc2, suppressed PI3K-Akt-mTOR, and phenocopied hyperoxia; Smarcc2 knockdown had opposite effects. Baf60c bound Smarcc2 but not PI3K. PMVEC-specific Baf60c overexpression attenuated pulmonary hypertension and right ventricular hypertrophy and partially improved alveolar and microvascular injury.

**CONCLUSIONS:** Hyperoxia-induced BPD-PH is associated with reduced Baf60c, increased Smarcc2, and suppressed PI3K-Akt-mTOR signaling in PMVECs. Baf60c may indirectly regulate this pathway through Smarcc2. Endothelial Baf60c is a potential therapeutic target in BPD-PH.

**Novelty and Relevance:** *What Is New?:* A neonatal hyperoxia BPD-PH model was characterized with adverse hemodynamic and pulmonary vascular endpoints together with PMVEC functional impairment. Hyperoxia downregulates Baf60c, upregulates Smarcc2, and suppresses PI3K-Akt-mTOR signaling in PMVECs. Baf60c interacts with Smarcc2 without direct PI3K binding. PMVEC-specific Baf60c overexpression via superficial facial vein AAV delivery ameliorates pulmonary hypertension.

*What Is Relevant?:* BPD-PH is a developmental pulmonary hypertension phenotype in which pulmonary vascular resistance and right ventricular load are measurable with RVSP and Fulton index, consistent with Group 3 PH biology. SWI/SNF-mediated chromatin remodeling in pulmonary endothelium may link neonatal oxygen injury to impaired angiogenic endothelial function.

*Clinical/Pathophysiological Implications?:* Endothelial Baf60c deficiency may contribute to pulmonary vascular rarefaction after neonatal hyperoxia. Restoring endothelial Baf60c may improve pulmonary hemodynamics in BPD-PH, although structural lung injury may be only partially reversible.

## Introduction

Pulmonary hypertension (PH) complicates bronchopulmonary dysplasia (BPD) in a substantial proportion of extremely preterm infants and is associated with right ventricular dysfunction, prolonged hospitalization, and increased mortality.^1–4^ Bronchopulmonary dysplasia–associated PH (BPD-PH) shares features with Group 3 PH, in which alveolar hypoxia and impaired vascular growth contribute to elevated pulmonary vascular resistance.^2,4,5^ Effective pharmacologic strategies for BPD-PH remain limited, and cell type–specific mechanisms coupling lung injury to pulmonary vascular dysfunction are incompletely understood.^2,3^

PMVECs regulate barrier function, proliferation, migration, invasion, and crosstalk with mural cells during vascular development.^6–9^ In hyperoxia-exposed neonatal lungs, endothelial dysfunction and capillary rarefaction accompany simplified alveolar architecture and elevated RVSP in rodent models.^10–12^ However, how neonatal oxygen injury is transmitted to PMVEC signaling failure—and whether chromatin remodeling factors participate—remains unclear.^13,14^

Baf60c (BRG1-associated factor 60c; gene Smarcd3) and Smarcc2 (BAF170) are SWI/SNF subunits implicated in vascular and chromatin regulation.^15–23^ The PI3K-Akt-mTOR cascade supports endothelial survival, motility, and angiogenesis.^6,24^ Whether reciprocal Baf60c–Smarcc2 dysregulation links hyperoxia to PI3K-Akt-mTOR suppression in BPD-PH has not been established.

We hypothesized that neonatal hyperoxia reduces endothelial Baf60c, increases Smarcc2, and thereby suppresses PI3K-Akt-mTOR signaling, leading to PMVEC dysfunction, pulmonary vascular remodeling, and elevated RVSP with right ventricular hypertrophy, restoring endothelial Baf60c would reactivate this pathway and ameliorate BPD-PH. Here, we conducted phenotypic characterization of a neonatal mouse model of BPD-PH, defined PMVEC functional and transcriptomic changes, delineated the Baf60c–Smarcc2–PI3K-Akt-mTOR axis, and tested PMVEC-specific Baf60c overexpression *in vivo*.

## Methods

### Data Availability

Lizhong Du had full access to all data in the study and takes responsibility for the integrity of the data and the accuracy of the data analysis. RNA-seq accession number will be deposited upon acceptance.

### Animals

All procedures were approved by the Animal Ethics and Use Committee of Zhejiang University (approval ZJU20230034) and performed in accordance with the Chinese national regulations and guidelines for laboratory animal welfare and ethics, including GB/T 35892-2018. C57BL/6 mice (Shanghai SLAC Laboratory Animal Co., Ltd.) were housed in the specific pathogen–free facility at Zhejiang Chinese Medical University (22°C–24°C, 50%–60% humidity, 12-hour light/dark cycle, ad libitum chow and water). Breeders were paired at 8 weeks of age; vaginal smears defined the mating day. Pups were randomized within 24 hours after birth to normoxia (~21% O_2_) or hyperoxia.^25^

### Neonatal Hyperoxia Model

Hyperoxia pups were placed in an Ox-100HE chamber at 85% FiO_2_ from postnatal day 0 (P0) through P14, with daily replacement of sodium lime and anhydrous CaCl_2_.^10–12^ Dams were exchanged between normoxia and hyperoxia cages every 24 hours to limit maternal exposure effects.^10^ Endpoints were assessed on P14 (Figure 1A). This protocol induces alveolar simplification and vascular growth arrest characteristic of experimental BPD.^10,11,26^

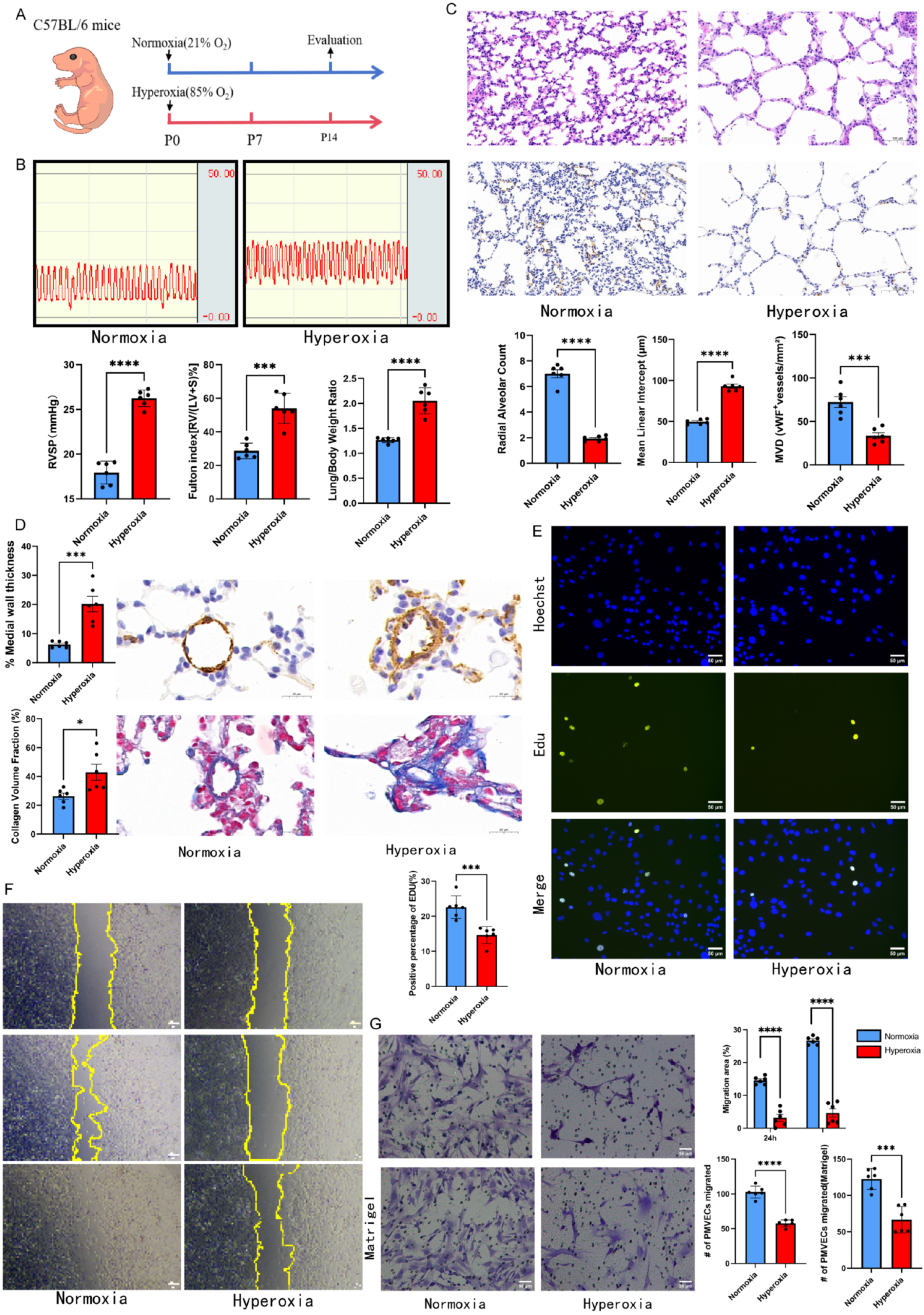

### Hemodynamic and Morphometric Measurements

RVSP was measured under pentobarbital (30 mg/kg, intraperitoneally) with ventilator support (DM-3000); a 26-gauge needle was inserted into the right ventricle, and mean RVSP was recorded with AcqKnowledge 4.0 (BIOPAC).^6,7^ Fulton index = RV/(LV+S); lung weight index = lung wet weight/body weight.^6^

### Histology and Immunostaining

Lungs were fixed in 4% paraformaldehyde, embedded in paraffin, and sectioned at 5 μm. HE, Masson, radial alveolar counts (RAC), mean linear intercept (MLI), α-smooth muscle actin immunohistochemistry (1:500), and von Willebrand factor immunohistochemistry (1:100) were performed as previously described.^7,10^ Arteriolar medial area percentage and vWF-positive vessels (10–50 μm) were quantified with ImageJ. Immunofluorescence for BAF60C, SMARCC2, and CD31 followed laboratory protocols.^16^

### PMVECs

PMVECs were isolated by explant adherence culture in ECGM supplemented with 5% serum and 90 U/mL heparin; passages 2–5 were used.^6–8^ CD31 positivity was >95% and α-smooth muscle actin negativity was confirmed (Supplementary Figure 1A).^6,7,9^

### Functional Assays

EdU incorporation (50 μM, 2 hours), Transwell migration (8-μm pore, 24 hours), scratch assay (0, 24, 48 hours), and Matrigel invasion (6.67%, 48 hours) were performed as described.^6,8^

### RNA-Seq and Bioinformatics

RNA was sequenced on NovaSeq 6000. Differential expression and pathway analyses used ClueGO/CluePedia in Cytoscape (Padj < 0.05).^6,27^

### qPCR, Western Blot, Co-IP, and LY294002

RNA and protein were extracted using standard protocols. qPCR used β-actin as internal control. Western blotting and coimmunoprecipitation with Protein A/G magnetic beads (MCE) followed published procedures,^6,16,20^ Co-IP was performed on PMVECs with Protein A/G magnetic beads using anti-BAF60C (5 μg), anti-SMARCC2 (3 μg), or control IgG (3 μg); complexes were rotated overnight at 4°C, captured on beads for 4 h, washed, eluted, and analyzed by Western blot (full protocol in Supplementary Methods S6). Pharmacologic inhibition: PMVECs were treated with the PI3K inhibitor LY294002 (50 μM) or DMSO vehicle for 24 hours to inhibit PI3K-Akt signaling.^6^ siRNA sequences and primers are in Supplemental Methods.

### Plasmid Transfection and siRNA

siRNA knockdown: Cells were transfected with siBaf60c or siSmarcc2 (sequences in Supplementary Table S4) using Lipofectamine RNAiMAX (30 pmol per well).Plasmid overexpression: The GV657-Baf60c construct was generated as described in Supplementary Methods S7 and Supplementary Figure 1E.^16,21^

### AAV *In Vivo*

LungX AAV carrying pAAV-TIE1p-Smarcd3-3×FLAG-P2A-mCherry-WPRE was used. Groups: normoxia control; hyperoxia + NC-AAV (1×10^11^ vg); hyperoxia + Baf60c-AAV (1×10^11^ vg). Under light isoflurane anesthesia, virus was injected via the superficial facial vein on day 1 of chamber exposure (P0–P1), as established for neonatal systemic AAV delivery.^28–30^

### Statistical Analysis

GraphPad Prism 9.0.0 and SPSS 27.0; mean±SEM. Two-group comparisons: two-tailed unpaired Student t test (or Mann-Whitney U if nonnormal). Multi-group experiments with two categorical factors (oxygen exposure × oeBaf6c treatment): two-way ANOVA with Tukey post hoc test (Figures 5C–5E). For Figure 6D and 6E, data from three groups (normoxia control, hyperoxia + NC-AAV, hyperoxia + Baf60c-AAV) were analyzed by one-way ANOVA followed by Tukey post hoc test. Figure 6B and 6C were analyzed as noted in figure legends. Figure 1 and two-group in vitro panels: n = 6 per group (animals or group comparisons in Figure 1E–1G) or n = 3 independent experiments, Student t test, as in figure legends. *P* < 0.05 was significant.

Detailed protocols are provided in the Supplemental Material (Tables S1–S5; Supplementary Methods S1–S9).

## Results

### 1. Establishment and phenotypic characterization of the BPD-PH mouse model

A BPD-PH mouse model was established as outlined in Figure 1A, using 85% FiO_2_ from P0 through P14.^10–12^ Compared with normoxia controls, the hyperoxia group showed markedly increased RVSP (Figure 1B, *P* < 0.0001), right ventricular hypertrophy index (Fulton index, RV/[LV+S]), right ventricular weight ratio (P < 0.001), and lung weight index (*P* < 0.0001). Hematoxylin–eosin staining revealed decreased radial alveolar counts and increased mean linear intercept (both P < 0.0001), indicating impaired alveolarization (Figure 1C). von Willebrand factor immunohistochemistry showed reduced pulmonary capillary density (*P* < 0.0001). α-Smooth muscle actin staining demonstrated medial thickening in pulmonary arterioles (10–50 μm) (*P* < 0.001), and Masson staining revealed increased collagen deposition (*P* < 0.05) (Figure 1D). Together, these data established a BPD-PH phenotype with pulmonary hypertension, right ventricular hypertrophy, pulmonary edema, impaired alveolar development, and pulmonary vascular remodeling.

### 2. Impaired proliferation, migration, and invasion of PMVECs in hyperoxia-exposed BPD-PH mice

Primary PMVECs were isolated from 14-day-old mice after hemodynamic measurement. Cultured cells showed cobblestone morphology, >95% were CD31-positive and α-SMA–negative (Supplementary Figure 1A and 1B). Compared with normoxia controls, PMVECs from hyperoxia-exposed mice showed reduced proliferation (Figure 1E, *P* < 0.001), migration (Figure 1F, *P* < 0.0001), and invasion (Figure 1G, *P* < 0.001). This *in vitro* phenotype paralleled reduced vWF-positive microvascular density in hyperoxia lungs (Figure 1D).

### 3. Hyperoxia exposure suppresses the PI3K-Akt signaling pathway in PMVECs

Transcriptome sequencing identified 1252 upregulated and 1817 downregulated genes in hyperoxia versus normoxia PMVECs (Supplementary Figure 1C). GO analysis showed enrichment of DNA binding, chromatin remodeling, cell signaling, motility, and adhesion (Figure 2A). KEGG analysis highlighted PI3K-Akt signaling (Figure 2B). Western blot confirmed suppressed PI3K-Akt activity in hyperoxia PMVECs (Figure 3A).

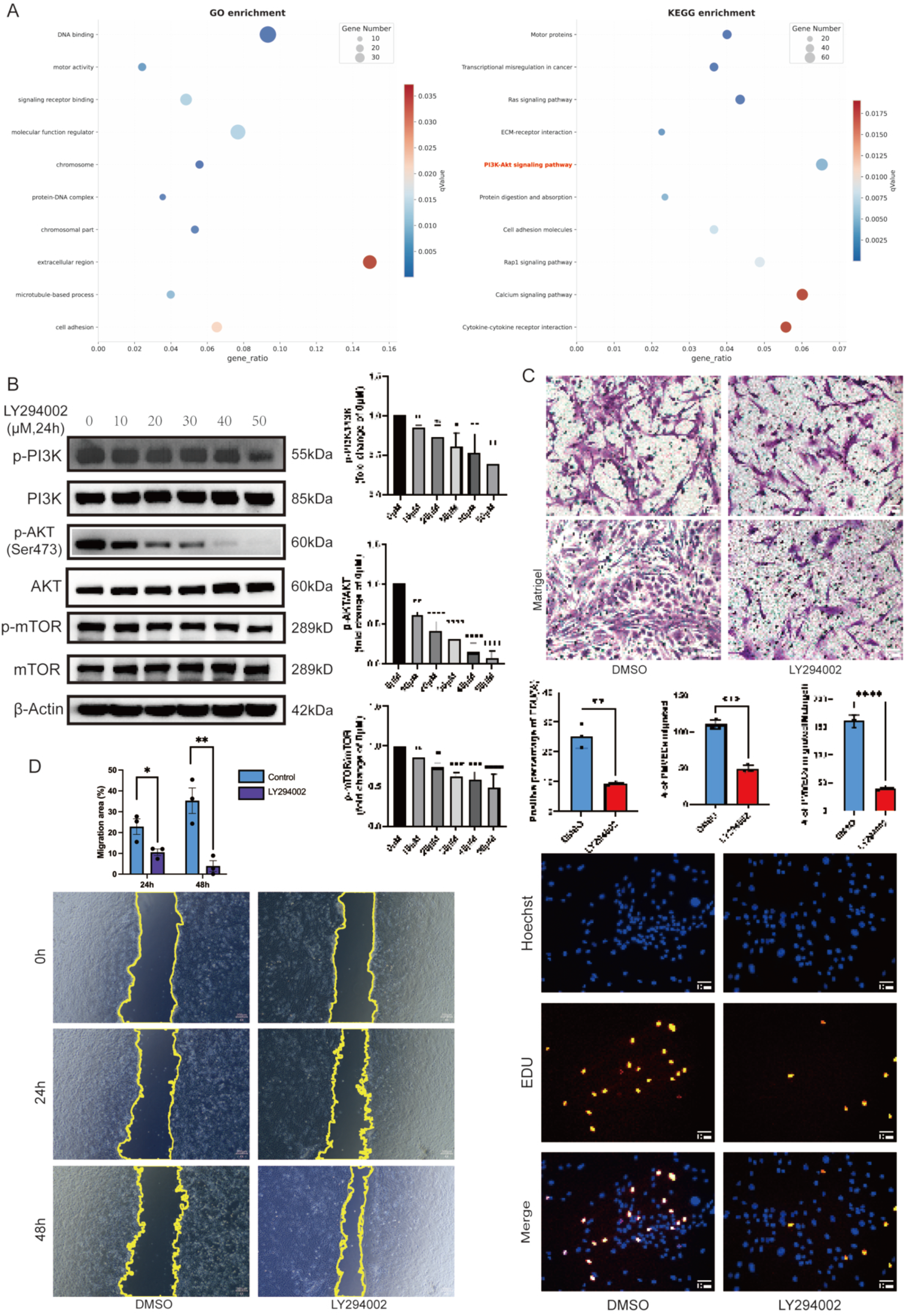

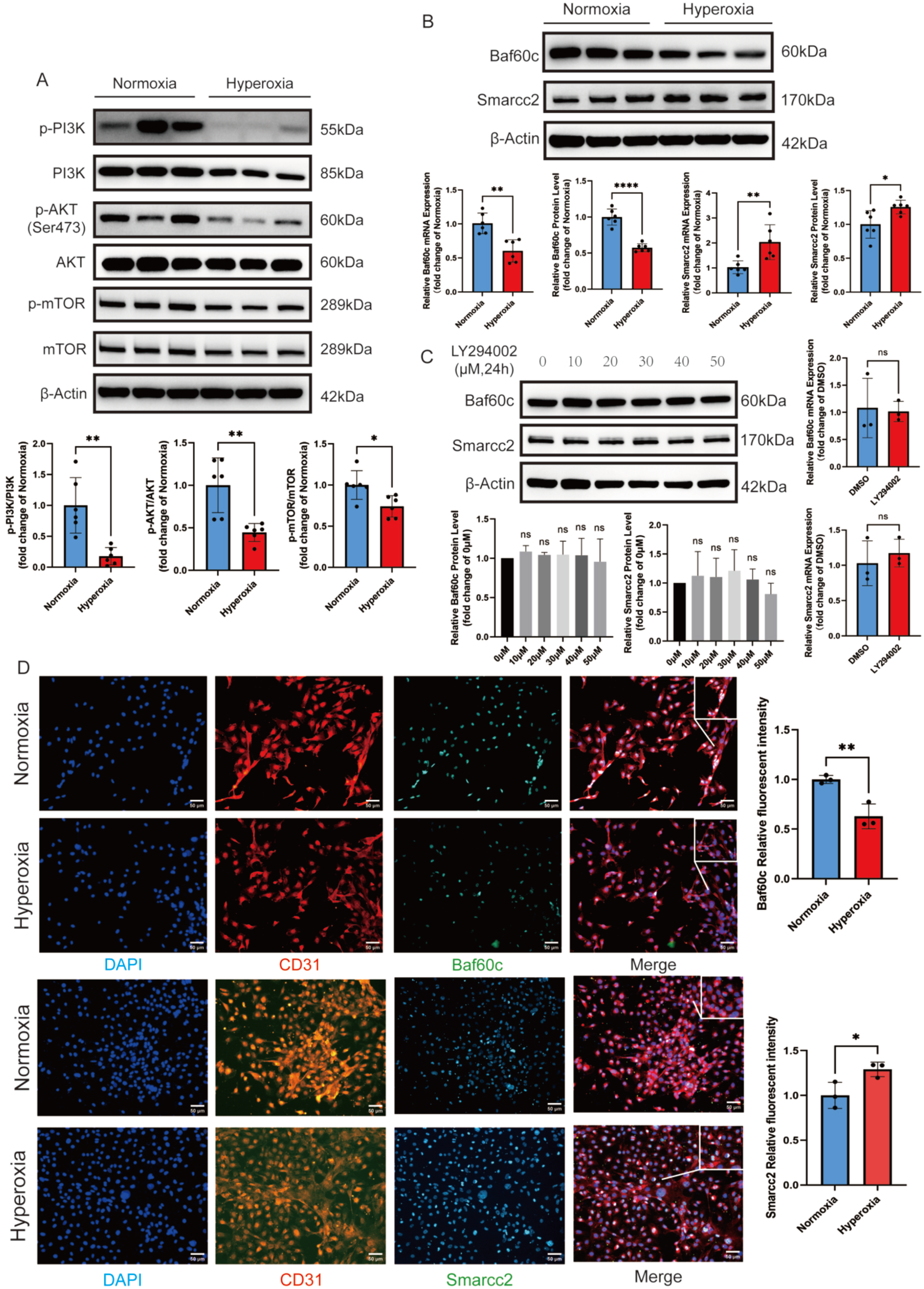

LY294002 inhibited PI3K-Akt in a dose-dependent manner (Figure 2C) and reduced PMVEC proliferation (*P* < 0.01), migration (*P* < 0.001), and invasion (*P* < 0.0001) (Figure 2D–2F), phenocopying hyperoxia-induced functional impairment.

### 4. Baf60c indirectly regulates the PI3K-Akt-mTOR pathway through Smarcc2

Hyperoxia PMVECs showed decreased Baf60c (mRNA *P* < 0.01, protein *P* < 0.0001) and increased Smarcc2 (mRNA P < 0.01, protein P < 0.05), confirmed by immunofluorescence (Figure 3B and 3D, Supplementary Figure 1D). PI3K-Akt activity was concurrently suppressed (Figure 3A). LY294002 did not alter Baf60c or Smarcc2 expression (Figure 3C).

Baf60c knockdown upregulated Smarcc2 and suppressed PI3K-Akt signaling (Figure 4A and 4B) and reduced PMVEC proliferation, migration, and invasion (Figure 4C–4E). Smarcc2 knockdown activated PI3K-Akt (Figure 4F) without changing Baf60c (Figure 4G). Co-immunoprecipitation showed Baf60c–Smarcc2 interaction but no direct binding to PI3K, after siBaf60c treatment, Smarcc2 expression and Baf60c–Smarcc2 association were further increased, whereas neither protein bound PI3K directly (Figure 4H).

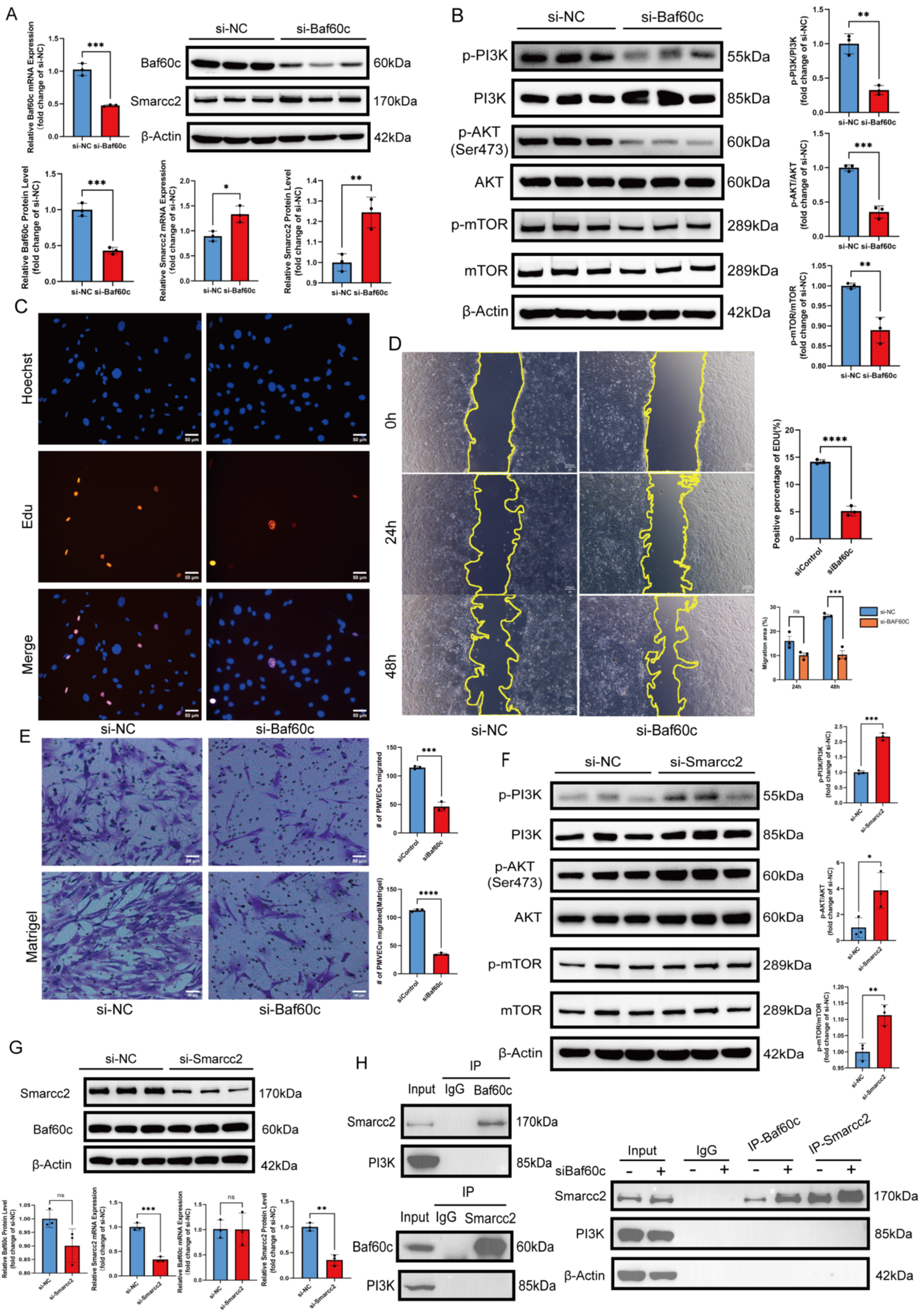

Smarcc2 knockdown enhanced PMVEC proliferation, migration, and invasion (Figure 5A–5C). Baf60c overexpression decreased Smarcc2, activated PI3K-Akt-mTOR (Figure 5D and 5E), and improved PMVEC function (Figure 5F–5H).

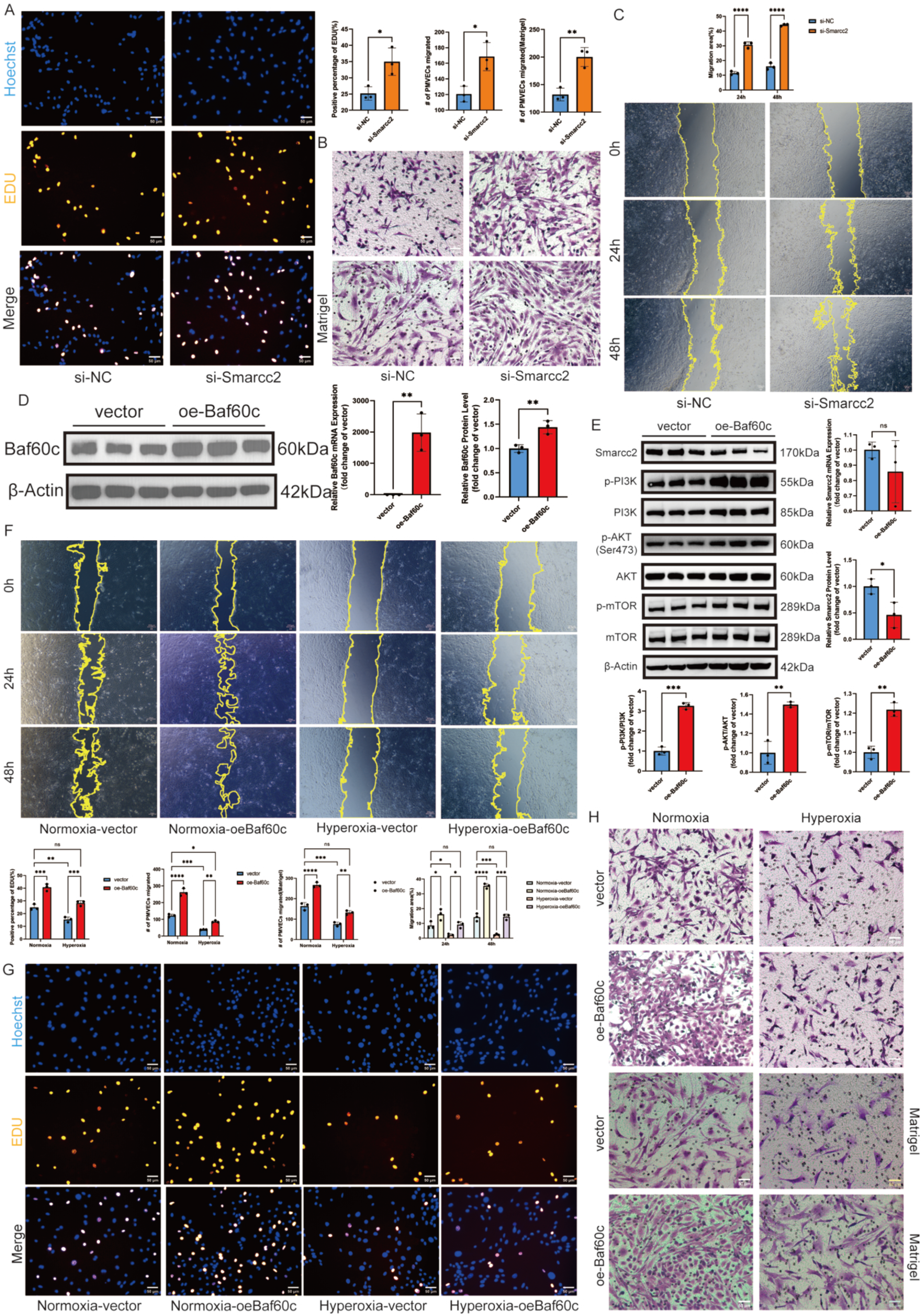

### 5. Endothelial-specific Baf60c overexpression ameliorates BPD-PH

Tie1-driven LungX AAV was injected via the superficial facial vein within24 hours after birth, followed by 14 days of 85% hyperoxia (Figure 6A). qPCR, Western blot, and immunofluorescence confirmed Baf60c overexpression in lung tissue and PMVECs (Figure 6B and 6C, Supplementary Figure 1G–1I). Baf60c-AAV significantly attenuated hyperoxia-induced elevations in RVSP (*P* < 0.0001), Fulton index (*P* < 0.01), and lung weight index (*P* < 0.001) (Figure 6D). Histology showed attenuated alveolar simplification (P < 0.001), reduced arteriolar medial and adventitial thickening (*P* < 0.05 and P < 0.001), and partial restoration of microvascular density (*P* < 0.01) (Figure 6E). Rescue of alveolar and microvascular abnormalities was partial.

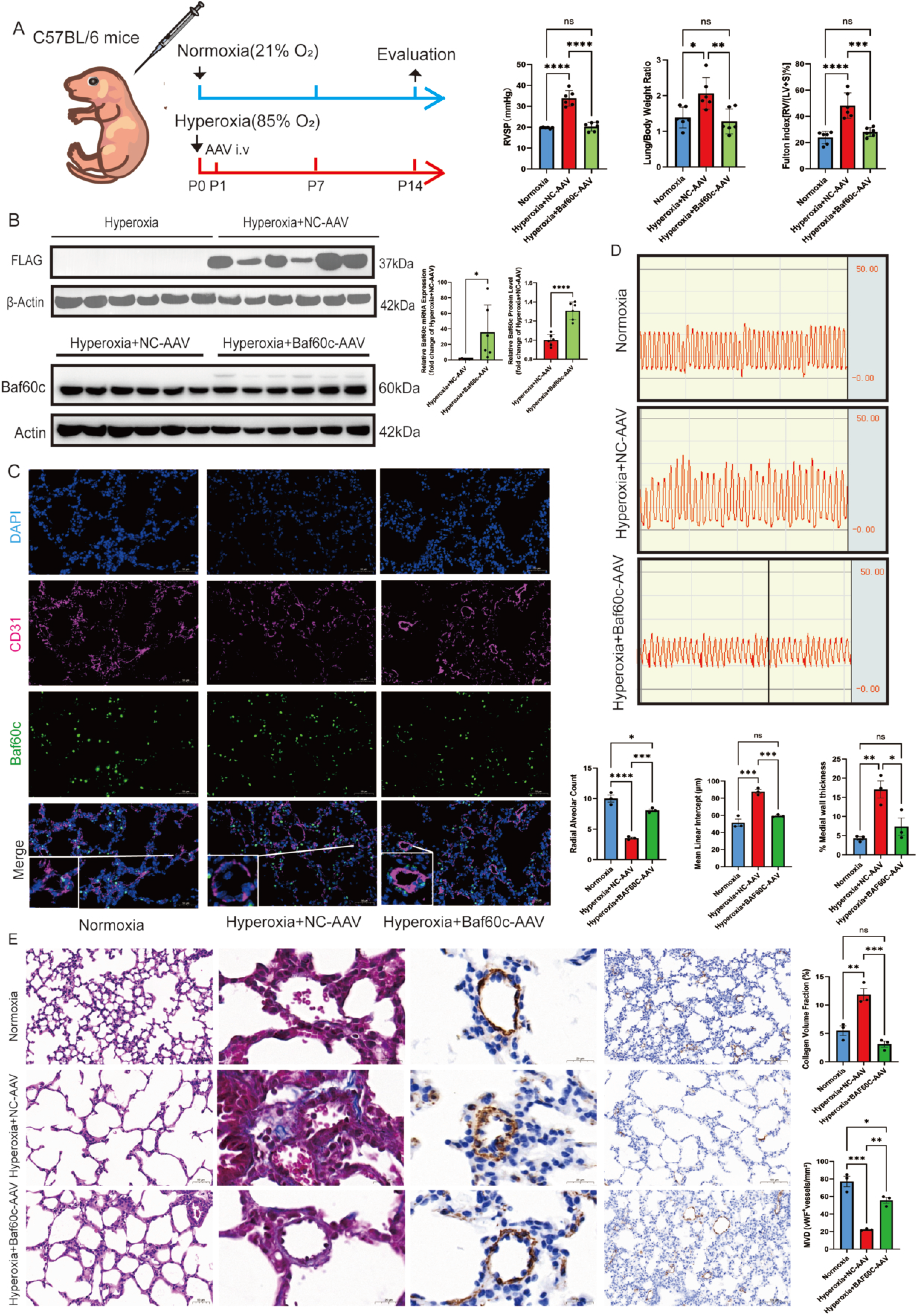

## Discussion

We report that neonatal hyperoxia produces a BPD-PH phenotype with elevated RVSP, right ventricular hypertrophy, pulmonary edema, impaired alveolarization, capillary rarefaction, and pulmonary arteriolar remodeling. At the cellular level, PMVECs show impaired proliferation, migration, and invasion associated with PI3K-Akt-mTOR suppression. Mechanistically, hyperoxia reciprocally regulates Baf60c and Smarcc2, Baf60c interacts with Smarcc2 without binding PI3K directly. PMVEC-targeted Baf60c restoration via superficial facial vein AAV delivery improves hemodynamics and partially reverses vascular structural injury. Together, these findings position endothelial Baf60c–Smarcc2–PI3K-Akt-mTOR signaling as a contributor to BPD-PH pathobiology and a potential therapeutic axis. The integrated mechanism is summarized in Supplementary Figure S2.

BPD-PH remains a major complication of prematurity, with measurable hemodynamic burden in pediatric cohorts followed over years.^1–4,31^ In preterm infants, early pulmonary vascular injury has been linked to impaired bone morphogenetic protein (BMP) signaling and other developmental risk factors, underscoring that the neonatal pulmonary circulation is vulnerable during lung maturation.^32^ Elevated pulmonary artery pressure in BPD-PH reflects reduced vascular cross-sectional area, adverse remodeling of small pulmonary arterioles, and impaired angiogenesis.^6,33,34^ Clinical and experimental work consistently shows endothelial dysfunction, capillary rarefaction, and disordered angiogenesis in hyperoxia-exposed neonatal lungs.^6,10–12,35^ In our model, medial thickening and increased collagen deposition parallel capillary loss-findings aligned with experimental BPD-PH in which pulmonary arteriolar remodeling and smooth muscle–related pathways contribute to right ventricular afterload.^34,36^

Our hyperoxia protocol (85% O2, P0–P14, dam rotation) follows widely used neonatal BPD models in which alveolar simplification and vascular growth arrest are prominent.^10–12,26^ Adding RVSP and Fulton index links developmental lung injury to pulmonary vascular resistance and right heart load, endpoints central to the readership of Hypertension.^5,6,9^ The simplified alveolar phenotype we observed is consistent with arrested alveolar development described in severe BPD and in hyperoxia rodent models.^3,10,11,26,35^ Transcriptomic enrichment of chromatin remodeling and DNA-binding processes after hyperoxia fits a broader literature,as well as our previous study^37^ showing that neonatal hyperoxia can leave persistent epigenetic marks in lung tissue, including histone signature disturbances at vascular-related loci such as eNOS^37^, NOS3 and STAT3.^14^ Integrative reviews similarly emphasize epigenetic–metabolic coupling as an underexplored layer in pulmonary vascular disease.^9,13,23,24,36,38^ In our data, GO enrichment and reciprocal Baf60c–Smarcc2 dysregulation suggest that oxygen injury may be transmitted partly through chromatin-level reprogramming in PMVECs, not only through acute hypoxic signaling. This extends prior hyperoxia epigenetics work by linking SWI/SNF subunits to endothelial PI3K-Akt dysfunction in BPD-PH. The concordance between KEGG enrichment, suppressed PI3K-Akt activity in hyperoxia PMVECs, and functional mimicry by LY294002 supports PI3K-Akt as a functionally relevant effector rather than a secondary transcriptional correlate. This aligns with endothelial PI3K-Akt biology in pulmonary vascular disease.^6,24,38^ Recent work in Hypertension showed that targeting an endothelial ENO1–PI3K-Akt-mTOR axis can alleviate hypoxic PH,^6^ supporting the plausibility of endothelial kinase pathways as intervention nodes—though our model emphasizes developmental hyperoxia rather than adult hypoxia alone. Importantly, LY294002 did not alter Baf60c or Smarcc2 expression, placing SWI/SNF-related regulation upstream of or parallel to PI3K rather than downstream of Akt activation.^16,20^

Loss- and gain-of-function data indicate that Baf60c and Smarcc2 act in opposition in PMVECs: Baf60c knockdown resembles hyperoxia, whereas Smarcc2 knockdown opposes it. Co-IP confirms physical association between Baf60c and Smarcc2 but not direct binding to PI3K, supporting indirect regulation of PI3K-Akt-mTOR. Baf60c knockdown not only increased Smarcc2 abundance but also enhanced their physical association, supporting reciprocal regulation within the SWI/SNF complex without direct recruitment of PI3K. This interpretation sits within established SWI/SNF biology. Baf60c prevents vascular smooth muscle de-differentiation and supports vascular homeostasis through epigenetic control.^16,17^ Independently, Smarcd3 has been identified as an epigenetic modulator reshaping transcriptional programs, supporting a regulatory rather than purely structural role for Baf60c.^18^ Smarcc2 participates in chromatin remodeling linked to proliferative signaling in other contexts.^20,21,39^ Structurally, Smarcc2 is a core scaffold of the PBAF SWI/SNF complex organized on nucleosomes.^19^ The Baf60c–Smarcc2 interaction we detect is compatible with coexistence within SWI/SNF/PBAF assemblies, as suggested by nucleosome-bound BAF/PBAF structural studies.^17,19^ KLF4-dependent SWI/SNF recruitment in endothelial enhancer reprogramming further supports chromatin remodeling as a gatekeeper of endothelial phenotype.^22^ Our data extend this framework from smooth muscle and generic endothelial shear responses to pulmonary microvascular endothelium after neonatal oxygen injury.

We propose that hyperoxia shifts the balance toward Smarcc2-dominant chromatin remodeling that suppresses a proangiogenic PI3K-Akt-mTOR program normally maintained by Baf60c. This model does not require direct Baf60c binding to PI3K, instead, Smarcc2 may remodel accessibility at promoters or enhancers of pathway components—a hypothesis for future chromatin profiling.

Tie1-driven Baf60c-AAV delivered by superficial facial vein injection at postnatal day 1 reduced RVSP, right ventricular hypertrophy, and lung weight gain after hyperoxia. Neonatal superficial facial vein delivery is an established route for systemic AAV at P0–P1 and enables intervention during rapid lung vascular development.^28–30^ As shown in Supplementary Figure 1G–I, Tie1-driven AAV transduced lung tissue and PMVECs, supporting endothelial-biased Baf60c rescue in vivo.

Hemodynamic improvement with only partial reversal of alveolar simplification suggests that endothelial rescue can reduce pulmonary vascular load even when parenchymal architecture remains incompletely restored.^2,3,34,36^ This pattern parallels gene or pathway restoration in developmental lung disease improving PH-related outcomes while leaving residual parenchymal injury.^40,41^ Endothelial-targeted STAT3 delivery, for example, alleviated PH in alveolar capillary dysplasia models while structural defects persisted.^40^ Similarly, interventions targeting neonatal lung growth axes can improve vascular outcomes without full normalization of alveolar structure.^26,41^ Our partial microvascular and arteriolar recovery is therefore biologically plausible: Baf60c restoration preferentially benefits the pulmonary circulation and microvasculature, whereas broad alveolar repair may require longer duration or combined strategies.^10–12,40^

### Limitations

This study used a single hyperoxia intensity and duration, one mouse strain, and primarily male pups unless otherwise balanced. We did not perform chromatin immunoprecipitation or assess genome-wide accessibility changes at PI3K-Akt loci. Clinical translation requires validation in large-animal models and human BPD-PH specimens. Future work should define optimal AAV timing, dose, and long-term safety, test combination with alveolarization-promoting strategies and integrate BMP pathway biology with chromatin remodeling given shared relevance to neonatal pulmonary vascular development.

## Conclusions

Neonatal hyperoxia-induced BPD-PH is associated with PMVEC dysfunction, reduced Baf60c, increased Smarcc2, and suppressed PI3K-Akt-mTOR signaling. Baf60c interacts with Smarcc2 and indirectly modulates PI3K-Akt-mTOR activity. PMVEC-specific Baf60c overexpression via superficial facial vein AAV delivery ameliorates pulmonary hypertension and partially improves pulmonary vascular structure. Targeting the endothelial Baf60c–Smarcc2 axis may offer a strategy for BPD-PH.

## Non-standard Abbreviations and Acronyms

AAV: adeno-associated virus
BPD: bronchopulmonary dysplasia
BPD-PH: bronchopulmonary dysplasia–associated pulmonary hypertension
Co-IP: coimmunoprecipitation
EdU: 5-ethynyl-2′-deoxyuridine
FiO_2_: fraction of inspired oxygen
HE: hematoxylin–eosin
MLI: mean linear intercept
NC-AAV: negative-control AAV
oeBaf60c: Baf60c overexpression
PH: pulmonary hypertension
PMVEC: pulmonary microvascular endothelial cell
RAC: radial alveolar count
RV: right ventricle
RVSP: right ventricular systolic pressure
SWI/SNF: switch/sucrose nonfermentable
vWF: von Willebrand factor

## Acknowledgement

We thank National Clinical Research Center for Child Health, Zhejiang Chinese Medical University Laboratory Animal Center for technical assistance.

## Author contributions

Q.L., L.Z., L.D. conceived the study and acquired funding. L.D. supervised experiments and co-wrote the manuscript. Q.L., L.Z., Q.C. established the BPD-PH model and performed hemodynamic and histologic phenotyping. Q.L., Q.W., L.Z., K.C.,C.H., isolated PMVECs. Q.L., Q.W. performed molecular biochemistry (knockdown/inhibitor assays, plasmid construction, qPCR, Western blot, Co-IP) and AAV-mediated in vivo rescue. Q.L., Q.C. performed statistics and image quantification. Q.L., L.D. drafted the manuscript. All authors edited and approved the final version.

